# The P3b differentiates parallel physical and rule-based updating of a sensory model

**DOI:** 10.1101/2021.02.17.431659

**Authors:** Michael Schwartze, Francesca I. Bellotti, Sonja A. Kotz

## Abstract

The capacity to form and update mental representations of the type and timing of sensory events is a central tenet of adaptive behavior in a dynamically changing environment. An internal model of stimulus contingencies provides a means to optimize behavior through predictive adjustments based on past to future events. To this end, neural and cognitive processes rely on systematic relations between events and use these rules to optimize information processing. The P3 complex of the event-related potential of the electroencephalogram (ERP/EEG) is a well-established and extensively tested index of such mechanisms. Here we investigated the P3b sensitivity to auditory stimulus deviations associated with two updating operations: physical change (switching stimulus pitches) and rule change (switching additive and subtractive target stimulus counting). Participants listened to a variant of the classical oddball sequence consisting of frequent standard (600 Hz) and two equally probable less frequent deviant tones (660 Hz, 540 Hz), keeping count of the deviant tones and switching between addition and subtraction with a pitch change. The results indicate specific amplitude modulations, confirming the P3b as a context-sensitive marker of physical and cognitive components of an internal model. This suggests that the P3b can be used as a differential marker of predictive coding mechanisms.

## 1. Introduction

Adequate reactivity and adaptation to change in the environment rank among the most fundamental tasks for any organism and are potentially facilitated by predictions about the future course of events, i.e., “adaptation to change by anticipation” (Fraisse, 1963). Across various processing levels, predictive adaptation to change may afford adequate and timely behaviour not only as a consequence but in active anticipation of sensory events to limit surprise and the expenditure of neural and cognitive resources that is associated with it. This notion is the central tenet of the seminal predictive coding framework, which posits that organisms thrive to reduce free energy (“surprise”) by adjusting models of the world towards the point of minimum prediction error by recognizing the most likely causes of the current sensory input (Friston, 2005; Clark, 2012; Friston, Thornton, & Clark 2012). In this framework, prediction error is understood as the difference between sensory input and predictions generated by the organism’s model of the environment. Stimulus events induce surprise to the extent to which they deviate from the model. Conversely, stimulus recognition is associated with the resolving of uncertainty, and consequently a reduction of surprise.

Sensory surprise is reflected in event-related responses of the electroencephalogram (ERP/EEG) such as the mismatch negativity and P3 (Friston, 2005; Garrido, Kilner, Stephan, & Friston, 2009; Chennu et al., 2013; Friston, Fitzgerald, Rigoli, Schwartenbeck, & Pezzulo, 2017; Heilbron & Chait, 2018). These ERPs can mirror the process by which a person infers causes from sensations on the basis of continuously updated models of the environment. The P3 is sensitive to the direction of attention and may indicate the need to optimize cognitive resource allocation to focal attending (Polich, 2007). The P3 is one of the most established functional markers in neurocognitive research (Sutton, Braren, Zubin, & John, 1965; Donchin, 1981; Linden, 2005; Polich, 2007). However, the classic P3 is an ERP complex that comprises at least two subcomponents. Separable in terms of neurochemical, distributional (scalp topography), temporal (peak latency), and functional characteristics, the P3a and P3b subcomponents are differentially indicative of attentional and decision-making processes (Linden, 2005; Polich & Criado, 2006; Polich, 2007). Typically evoked using “oddball-type” experimental paradigms that involve the presentation of infrequent deviant stimulus events in a train of repetitive and more frequently presented standard events, both subcomponents are markers of mechanisms that are related to the processing of changes in the configuration of the sensory environment. In particular the attention-dependent P3b has long been established as an electrophysiological marker of uncertainty and surprise (Sutton, Braren, Zubin, & John, 1965; Donchin, 1981), making it a versatile tool for the study of predictive adaptation to change by anticipation as a form of continuous adjustment to the environment towards the point of minimum prediction error.

The P3b is associated with the memorability of sensory events and a process of schema revision, i.e., an adaptation of a cognitive representation of the environment, with larger amplitudes indicating more surprising events (Donchin, 1981; Donchin & Coles, 1988). However, this process of schema revision critically depends on task-relevance and subjective probability (e.g., as defined by task-relevance) rather than on the objective relative probability of events (e.g., the actual ratio of expected and surprising events). In other words, it relies on predictions concerning the future course of events, with “task-relevance” defined as the ability of a stimulus to resolve uncertainties on the side of the participant, which in turn depends in part on what the participant attempts to do at a given time (Donchin 1981). These predictions are “active expectations” that draw on limited cognitive resources such as attention to balance readiness for events and the ability to respond to unexpected events (Kahnemann & Tversky, 1982). This implies the establishment of a subjective cognitive frame that involves the continuous maintenance of attention and task instructions in reference to an internal model of the stimulus environment. Enhanced P3b amplitudes for more surprising but also for predicted events suggest stronger induction and potentially also more efficient updating of the model (Miniussi, Wilding, Coull, & Nobre, 1999; Schwartze, Rothermich, Schmidt-Kassow, & Kotz, 2011). Critically, in this aspect and the associated amplitude enhancement effects, the P3b differs from earlier “exogenous” auditory ERP components such as the P50 or N1, which show suppression effects, i.e., attenuated amplitudes with temporal and formal (pitch) predictability (Lange 2009; Schwartze, Farrugia, & Kotz, 2013; for a review see Bendixen, SanMiguel, & Schröger, 2012). The amplitude enhancement of the P3b for surprising sensory events suggests that physical and task-related characteristics can have a differential impact on P3b amplitude, reflecting the extent of the subjective surprise that is associated with each of these aspects. The magnitude and functional differentiation of the P3b surprise response can therefore provide insight into the nature of the underlying model and cognitive resource allocation, aspects typically outside the focus of formal approaches to predictive coding. Like other electrophysiological correlates, this makes the P3b a surprise signal that carries specific information about *how* sensory data are surprising, i.e., about how input deviates from predictions (den Ouden, Kok, & de Lange, 2012).

The present study investigated whether and how the P3b would be differentially modulated by the context-defined role of physically identical deviant events in a continuous stimulus sequence. The primary goal was to assess the specificity and differentiation of the P3b in terms of a sensory target-model and cognitive-rule updating using a continuous three-tone oddball sequence (consisting of frequent standard and infrequent deviants of either higher or lower pitch). We hypothesized that P3b amplitude morphology would reflect the qualitative differences and the specific surprise associated with the respective type of schema revision. Task instructions introduced a distinction between (physical) “model-updating” (standard vs. target pitch) and “rule-updating” (deviant vs. other deviant pitch) by means of alternating additive and subtractive counting of the deviant tones. Participants were asked to continuously either add to or subtract from their count as long as a deviant was of the same type as the previous deviant and switched from addition to subtraction (and vice versa) if the deviant was of a different type. With this setup, we expected to observe larger P3b amplitudes for rule-updating due to an additional surprise component relative to model-updating alone. Thus, a differentiation of the surprise response indexed by the P3b could indicate independent aspects of the predictive adaptation to change in the environment.

## 2. Methods

### 2.1 Participants

Thirty-two healthy, right-handed volunteers (24 female, 8 male) participated in the study. They all reported normal hearing and no history of a neurological disorder. Age ranged from 19 to 28 years (M = 23.12, SD = 2.68). All participants gave their informed written consent and received either a compensatory fee or course credits. The study was approved by the ethical committee at the Faculty of Psychology and Neuroscience at Maastricht University (ECP-161_04_02_2016).

### 2.2 Experimental setup

Participants sat in an electrically-shielded sound-attenuated booth in front of a computer screen that continuously displayed an asterisk throughout the EEG recording session. Participants were asked to fixate the asterisk and to pay attention to a tone sequence that was presented via loudspeakers placed at a distance of approximately 1.5 m left and right of the screen. The stimulus sequence consisted of equidurational (150 ms) tones of three different pitches, frequent 600 Hz standard (N=256) and equiprobable infrequent 540 and 660 Hz deviant (N=32 each) tones that were presented with an stimulus-onset-asynchrony (SOA) of 1000 ms, and a corresponding inter-stimulus-interval (ISI) of 850 ms. Pseudo-randomization ensured that no more than two deviants were presented in a row. Participants were asked to refrain from any motor behavior and to perform a counting task (Fig.1). Starting with a value of 20, the task was to continuously add or subtract 1 if a deviant was of the same type as the preceding deviant (model update), and to switch to addition or subtraction if the deviant was of a different type as the preceding deviant (rule update).

**Fig 1.**
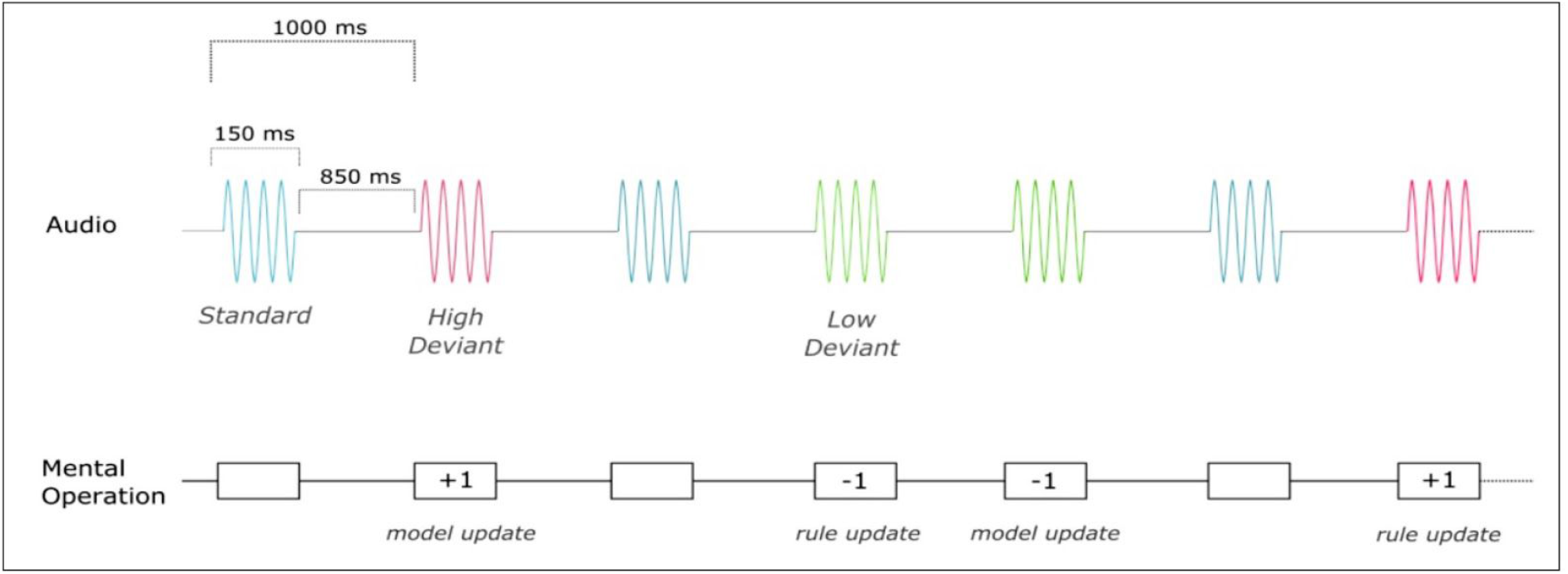
Stimulus sequences and experimental task. Infrequent 660 Hz and 540 Hz (pitch) deviant tones were presented among frequent 600 Hz standard tones. Participants counted deviant tones, adding or subtracting 1 to or from their count for each deviant of the same type and switching from addition to subtraction with each deviant of the other type. Deviants preceded by a deviant of the same type required an update of the physical stimulus model, whereas deviants preceded by a deviant of the other kind required an additional rule update.

### 2.3 EEG recordings and analyses

Continuous EEG was recorded with a sampling rate of 1000 Hz from 59 scalp-electrodes mounted into an elastic cap according to the 10-20 international system. Additional electrodes were placed on the sternum (ground) and on the left and right mastoid (for online reference and later re-referencing). Parallel horizontal and vertical electrooculography was recorded via electrodes placed on the outer canthus of both eyes and above and below the right eye, respectively. Stimulus randomization and presentation were controlled by Presentation 20 (Neurobehavioral Systems). During the recordings, electrode impedance was kept below 10 kΩ.

The Letswave6 toolbox (https://www.letswave.org) running on Matlab (Mathworks) was used for data processing. Data were bandpass-filtered using a Butterworth filter from 0.5 to 30 Hz (order 4). A combination of independent component analysis (ICA, Runica algorithm) and amplitude-based epoch rejection was applied to clean the data. Components dominated by eye-blinks were identified via ICA and removed from the data. The remaining data were then segmented into epochs lasting from -100 ms to 700 ms post-stimulus onset. Epochs exceeding ±100 μV at any electrode location were automatically rejected. Finally, some epochs containing residual artefacts, primarily slow drifts, were manually removed. Epochs were then averaged for each participant and then across all participants. Two regions of interest (ROIs) were defined for the analyses to verify and focus on the well-established centro-posterior scalp distribution of the P3b. To this end, one ROI comprising fronto-central electrodes (locations AF3, AFz, AF4, F3, F1, Fz, F2, F4, FC3, FC1, FCz, FC2, FC4) was contrasted with another ROI comprising centro-posterior electrodes (locations CP3, CP1, CPz, CP2, CP4, P3, P1, Pz, P2, P4, PO3, POz, PO4). Statistical analyses were performed in SPSS 25 (IBM). Initial analyses aimed at verifying the presence of a significant positive peak deflection around 300 ms in the averaged response across all channels. All analyses were performed on data derived from a time-window lasting from 290 to 450 ms relative to stimulus onset. The subsequent main analysis on re-grouped data was conducted using a 2 x 2 ANOVA that included the factors *region* (fronto-central vs. centro-posterior ROI) and *condition* (model-vs. rule-update) on mean ERP amplitudes.

## 3. Results

The values reported for the counting task (M = 22.69, SD = 2.88) indicated that participants understood and followed the instructions. Initial visual inspection of the ERP data confirmed the presence of the expected dominant positive deflections in response to both types of deviant tones relative to standard tones (Fig. 2).

**Fig 2.**
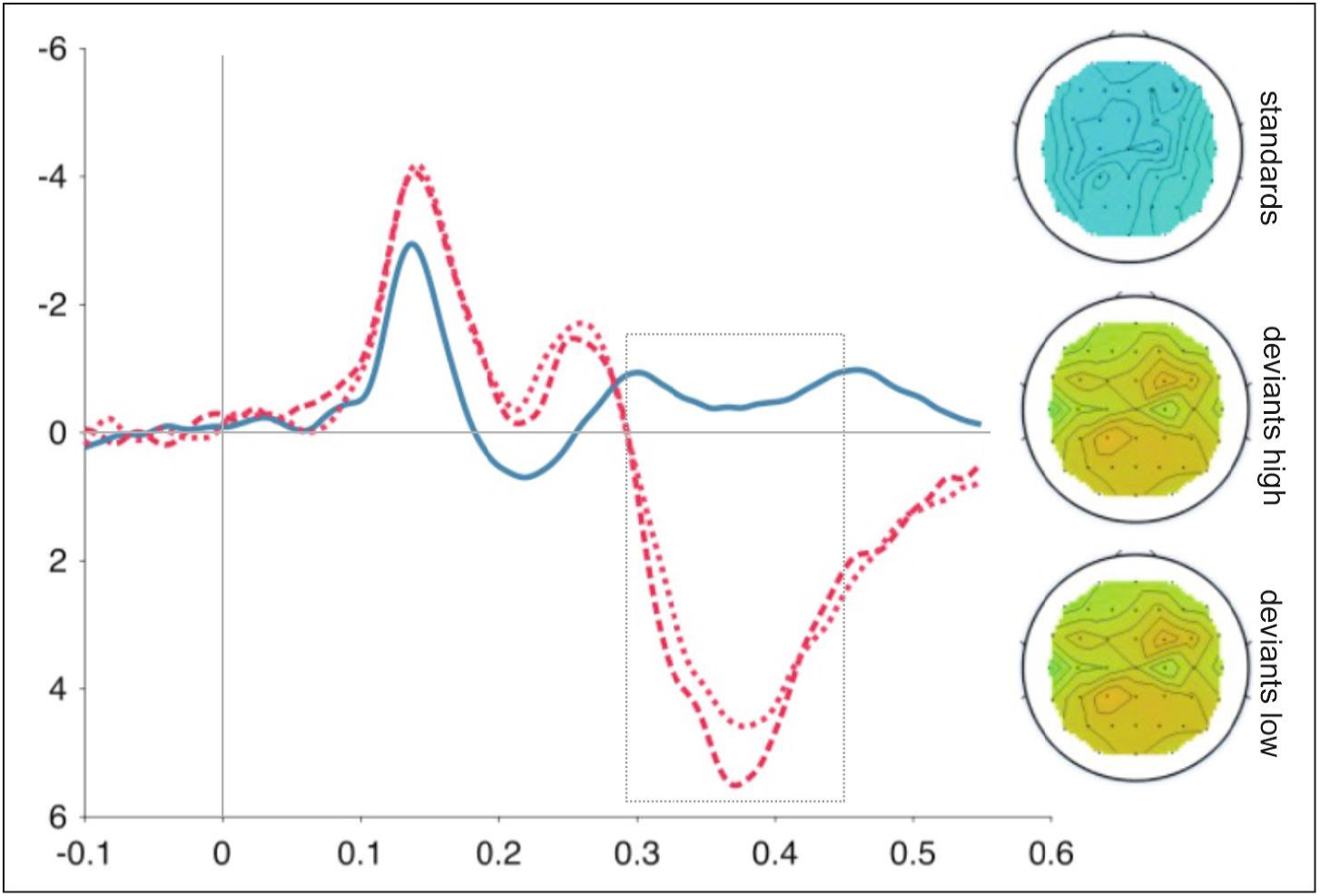
Event-related potential (ERP) results. Grand average ERP waveforms in response to standard (600 Hz, blue), high-pitch deviant (660 Hz, dashed red) and low-pitch deviant (540 Hz, dotted red) tones with corresponding scalp topographies (290-450 ms).

### 3.1 Peak deviance response

Paired samples t-tests were conducted to assess if the positive peaks differed from responses to standard tones. As expected, differences in peak amplitudes were found to be highly significant between standards (M = .41, SD = 1.36) and high deviants (M = 7.02, SD = 4.06); *t*(31) = -9.51, *p* < .01, *Cohen’s d*_*z*_ = -1.68 and between standards and low deviants (M = 6.36, SD = 3.43) *t*(31) = -11.51, *p* < .01, *d*_*z*_= -2.04. Notably, there was no significant difference in peak amplitudes when comparing low and high deviants; *t*(31) = -1.43, *p* = .16, *d*_*z*_ = -.25. These results confirmed that both deviants successfully elicited the expected response.

### 3.2 Differential rule and model updating

For the purpose of the subsequent main analysis, epochs for high and low deviants were regrouped according to their context-defined role in the sequence (Fig. 3). Deviants were pooled into one group whenever they were preceded by a deviant of the same type (model-update, i.e., requiring a physical/pitch update in response to the deviant relative to preceding standard tones) and into another group whenever they were preceded by a deviant of a different kind (rule-update, i.e., requiring a physical/pitch plus a counting rule update). Although the preceding analysis of the deviance response did not identify a significant difference between the two deviant responses, this procedure was adopted to ensure that subsequent findings could be attributed to context-defined roles rather than to physical stimulus differences.

**Fig 3.**
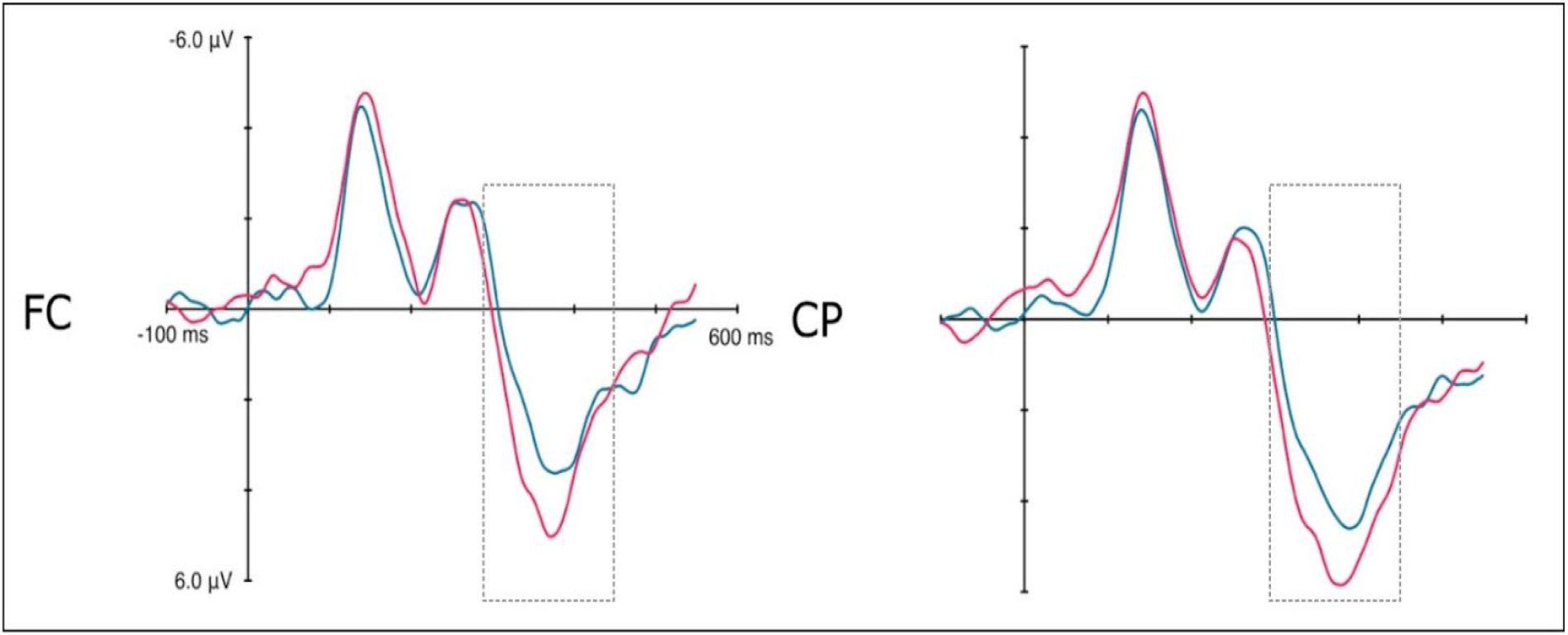
P3 differentiation as a function of model- and rule-updating. Grand average waveforms of responses recorded from fronto-central (left) and centro-posterior (right) regions associated with model-updating (blue) and rule-updating (red) deviant stimuli.

The results of the respective ANOVA showed a significant main effect of *condition, F*(1,31) = 18.15, *p* < .01. There was no main effect of *region* but a significant interaction of the factors *region* and *condition F*(1,31) = 4.95, *p* < .04. Resolving this interaction by the factor *region* to directly compare conditions revealed the hypothesized significant differences in the fronto-central *t*(31) = -3.06, *p* < .01, *d*_*z*_ = -.54 and the centro-posterior ROI *t*(31) = -4.89, *p* < .001, *d*_*z*_= -.86. These findings confirm the hypothesized differential sensitivity of the P3b to model- and rule-updating as implemented in the parallel oddball design.

## 4. Discussion

The current study investigated the P3b amplitude modulation associated with model- and rule-updating, i.e., two processes that are considered to reflect surprise reactions in response to an unexpected change in the sensory environment. The experimental setup targeted these processes by means of parallel changes of the auditory target pitch and of the task-setting in a continuous oddball design. We hypothesized that the P3b amplitude would differentiate between “rule-updating” (as evoked in response to the first deviant of a kind) and “model-updating” (as evoked in response to the second and subsequent deviants of a kind). The underlying principle and relevance of an associative link between changing physical stimulus characteristics and rule-based adaptive behaviour may be exemplified by the changing phases of a traffic light. In this visual example, a change of color may cue the slowing or initiation of movement as different forms of rule-based behaviour. An efficient internal model of the environment would require representations of both aspects and the monitoring of changes to each in order to guide behaviour. Smaller amplitudes for model- as compared to rule-updating confirmed that the P3b is a sensitive differential marker of these processes. Considering the prominent status of the P3b as a widely-applied marker of neurocognitive function (Linden, 2005; Polich, 2007), the current findings provide further insight into the basic cognitive mechanisms that underlie predictive adaptation of an organism to a dynamic sensory environment.

The context-updating theory of the P3b posits that the updating of an internal model of the environment is indexed by this component or potentially by a whole group of P3-like ERP responses that reflect various updating operations (Donchin & Coles, 1988; Polich, 2007; Brydges & Barcelo, 2018). Different updating operations may relate to stimulus detection, attentional or memory resource-allocation, decision-making, task-switching, or response requirements. The current experimental setup decidedly focussed on sensory processes, and did not require any overt response. In this aspect, the study differs from previous studies that investigated task preparation and execution and identified additive contributions to P3 amplitude based on cue- and task-updating (Perianez & Barcelo, 2009). Although the sensory focus still implies various updating operations, it makes the results directly relevant for the differentiation of aspects that combine into a predictive internal model of the sensory environment. Importantly, with the current experimental setup, we did not observe a significant difference in ERPs elicited by high-compared to low-pitched deviants. Such a difference in P3b amplitudes only emerged when the deviant stimuli were grouped according to their respective role in the sequence.

The notion of an internal model of the sensory environment as part of a hierarchy of models is also central to the predictive coding framework of brain function, which has become a leading theory in auditory neuroscience (Heilbron & Chait, 2018; Denham & Winkler, 2020). The formal predictive coding approach and the notion of an internal or “mental” model and model-updating nevertheless remain difficult to reconcile as many characteristics of the cognitive model, e.g., constraints posed by limited attentional or memory resources, remain elusive. However, the P3b amplitude has been shown to be sensitive to the degree of likelihood certainty as specified by the Bayesian framework, i.e., it indicates the probability for the evidence of a hypothesis about the nature of sensory input (Kolossa, Fingscheidt, Wessel, & Kopp, 2013; Kopp et al., 2016). The current finding in terms of physical model-updating could be explained on this basis, whereas the additional rule-updating result may hint at a qualitatively different cognitive updating operation that is solely defined by the cognitive task context.

Taken together, the current results show that the P3b component reflects deviation from a predictive model of the environment and different associated cognitive processes. It therefore provides a marker to study the role, relative contribution, and the interaction of physical and cognitive aspects as components of predictive coding and adaptation to the environment by anticipation.

